# The natural selection of metabolism and mass selects lifeforms from viruses to multicellular animals

**DOI:** 10.1101/087650

**Authors:** Lars Witting

**Affiliations:** Greenland Institute of Natural Resources, Box 570, DK-3900 Nuuk, Greenland

**Keywords:** Evolution, metabolism, body mass, life history, allometry, major transition, lifeform

## Abstract

I show that the natural selection of metabolism and mass is selecting for the major life history and allometric transitions that define lifeforms from viruses, over prokaryotes and larger unicells, to multicellular animals with sexual reproduction.

The proposed selection is driven by a mass specific metabolism that is selected as the pace of the resource handling that generates net energy for self-replication. This implies that an initial selection of mass is given by a dependence of mass specific metabolism on mass in replicators that are close to a lower size limit. A maximum dependence that is sublinear is shown to select for virus-like replicators with no intrinsic metabolism, no cell, and practically no mass. A maximum superlinear dependence is instead selecting for prokaryote-like self-replicating cells with asexual reproduction and incomplete metabolic pathways. These self-replicating cells have selection for increased net energy, and this generates a gradual unfolding of a population dynamic feed-back selection from interactive competition. The incomplete feed-back is shown to select for larger unicells with more developed metabolic pathways, and the completely developed feed-back to select for multicellular animals with sexual reproduction.

This model unifies natural selection from viruses to multicellular animals, and it provides a parsimonious explanation where allometries and major life history transitions evolve from the natural selection of metabolism and mass.

## 1 Introduction

With this study I aim for the first natural selection theory that is sufficient to explain the joint evolution of metabolism, body mass, major life-history transitions, and inter-specific allometries within and across lifeforms from viruses over prokaryotes and larger unicells to multicellular animals. The theory describes how mobile lifeforms with elevated metabolic rates and large body masses are selected as a deterministic consequence of the origin of replicating molecules, and I test the selection by its ability to explain transitions in the life history and allometric exponents across the tree of life.

The theory is proposed here in a joint study (Witting 2016a,b,c) that integrates primary selection on mass specific metabolism into the selection theory of Malthusian Relativity (Witting 1997, 2008). The original version of the theory explains the evolution of major life history transitions (Witting 1997, 2002a, 2007) and allometric exponents in large animals (Witting 1995) from the density dependent interactive competition that selects for large body masses.

Witting (2016a) extended the allometric component of the theory with primary selection on metabolism, and showed that mass specific metabolism can be selected as the pace of the resource handling that generates net energy for self-replication. This net energy generates population growth and interactive competition that selects for larger body masses, with the joint selection on metabolism and mass selecting for a range of inter-specific allometries that are observed in taxa from prokaryotes over larger unicells to multicellular animals (Witting 2016a).

Dependent upon the importance of metabolism for the selection of mass, the allometric exponent for mass specific metabolism is predicted to change from 3/4 over zero to −1/4 for interactive behaviour in two spatial dimensions (2D), and from 5/6 to −1/6 for three dimensional behaviour (3D, Witting 2016a). And with an average exponent around 0.84 (DeLong et al. 2010), it is suggested that the body mass variation in prokaryotesis selected from primary variation in mass specific metabolism. This contrasts to multicellular animals where exponents around −1/4 and −1/6 (Peters 1983; Witting 1995; Savage et al. 2004; McNab 2008; Capellini et al. 2010) indicate body mass variationthat is selected from primary variation in the handling and/or densities of the underlying resources. And with an apparent decline in the exponent with an increase in mass from about 0:61 over zero to −0.20 (Witting 2016a), the selection of mass in protozoa is suggested to shift more or less continuously from the mechanism that operates in prokaryotes to the mechanism that operates in multicellular animals.

While Witting (2016a) predicted this range of allometries, he did not predict the actual lifeforms of prokaryotes, larger unicells and multicellular animals. This would require a theory where the origin of replicating molecules selects for an increase in metabolism and mass with a succession of evolutionary transitions that resemble the transitions in the life history and allometric exponents that are observed between virus, prokaryotes, larger unicells, and multicellular animals. It is this evolution that is studied in the current paper, where the transitions between the four lifeforms are predicted as a directional succession that follows from primary selection on mass specific metabolism and mass.

As the proposed natural selection will explain the evolution of complete life histories from a few basic principles, it deals with the simultaneous selection of a diverse set of life history traits that include metabolism, mass, allometric exponents, reproduction, survival, abundance, home range, cellularity, senescence, and sexual reproduction. The joint theory has as a consequence of this many interrelated equations, and it may therefore appear to be unnecessary complex at first. But it is in fact extremely parsimonious because all the major transitions in the life history and the allometric exponents are selected by the primary selection that is necessary for the evolution of metabolism and mass.

The complete evolutionary unfolding from replicating molecules to multicellular animals with sexual reproduction is, in essence, explained by four simple principles. These are *i*) natural selection by differential self-replication in populations, *ii*) conservation of energy, *iii*) a mass specific metabolism that depends on mass in self-replicators that are close to a lower size limit, and *iv*) a gradual unfolding of density dependent interactive competition from the population growth of self-replication. The selection of virus and prokaryote like replicators requires only the first three principles, but the selection of larger unicells and multicellular animals is dependent on the gradual unfolding of the feed-back selection that follows from density dependent interactive competition.

## 2 The proposed selection

In order to have a single model that will predict lifeforms from viruses to multicellular animals I make the average life history in the population evolve in response to primary selection on metabolism and mass. The most basic selection is a mass specific metabolism that is selected as the pace of the resource handling that generates net energy for self-replication (Witting 2016a).

The selection of mass is then following from the increase in net energy, either by a dependence of metabolism on mass in self-replicators that are close to a lower size limit, and/or by the density dependent interactive competition that is generated by the population growth that follows from the average net energy in the population. The associated transitions in the interactive selection of mass is selecting for major life history transitions (see Witting 2002a for details), and body mass allometries are selected as a response to the primary changes in metabolism and mass by two processes that I refer to as the metabolic-rescaling and mass-rescaling selection of the life history (see Witting 2016a for details).

The current paper is a direct extension of Witting (2002a, 2016a), and I assume that the reader is familiar with both papers. All the parameters and basic selection relations that I use are defined in Witting (2016a, see Table 5 for parameter definitions), and Witting (2002a) is essential for the selection of major life history transitions. Witting (2016a) describes also the majority of the physiological and ecological relations that select on metabolism and mass, and it solves these equations for the potential range of allometric correlations. This shows how the selection of optimal density regulation and physiological trade-offs link selection on the home range and population density to selection on metabolism, time, net energy and mass; but it does not describe the specific selection of metabolism and mass that will make the equations unfold into species distributions with different body mass allometries for the different lifeforms.

The latter selection is studied in the sections below, with evolutionary transitions in the allometric exponents following from the gradual unfolding of the population dynamic feed-back selection that is imposed by density dependent interactive competition. This gradual unfolding creates a gradual increase in the resource bias across the intra-specific variation in competitive quality; a change in bias that is selecting, not only for an increase in mass, but also for life history transitions (Witting 1997, 2002a, 2007, 2008). And when these predictions are combined, we obtain the evolution of four major lifeforms that resemble those of viruses, prokaryotes, larger unicells, and multicellular animals.

### 2.1 Life history model

In this sub-section I give a brief introduction to the common life history that I use as the basis for the overall selection model. This model

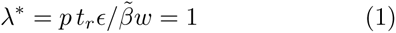

is a discrete formulation of the per generation replication (λ* = *pR* = 1 with *r*\* = ln λ = 0) of an average variant in an age-structured population that is situated at the dynamic equilibrium (*), where *p* is the probability to survive to reproduce, 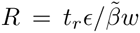 is lifetime reproduction, *t*_*r*_ the reproductive period in physical time, *ε* the net energy that is available for self-replication per unit physical time, *ω* the body mass, and 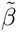 a scaling parameter that accounts for the energy that is metabolised by the offspring (see Witting 2016a for details).

The net energy for self-replication is a product

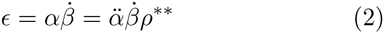

between the biochemical/mechanical handling of the resource (*α*) and the pace of this process (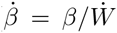), with pace being selected as mass specific metabolism (*β* with 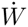 being the mass specific work of one joule metabolised per gram of tissue, see Witting 2016a for detail), and handling 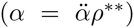 being defined for an implicit resource (*ρ*) at the density (*n*) regulated [*ρ* = *ρ*_*u*_*f*(*n*)] population dynamic equilibrium of the relevant selection attractor (**) for mass (Table 1), with 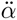 being the intrinsic component of resource handling.

**Table 1:**
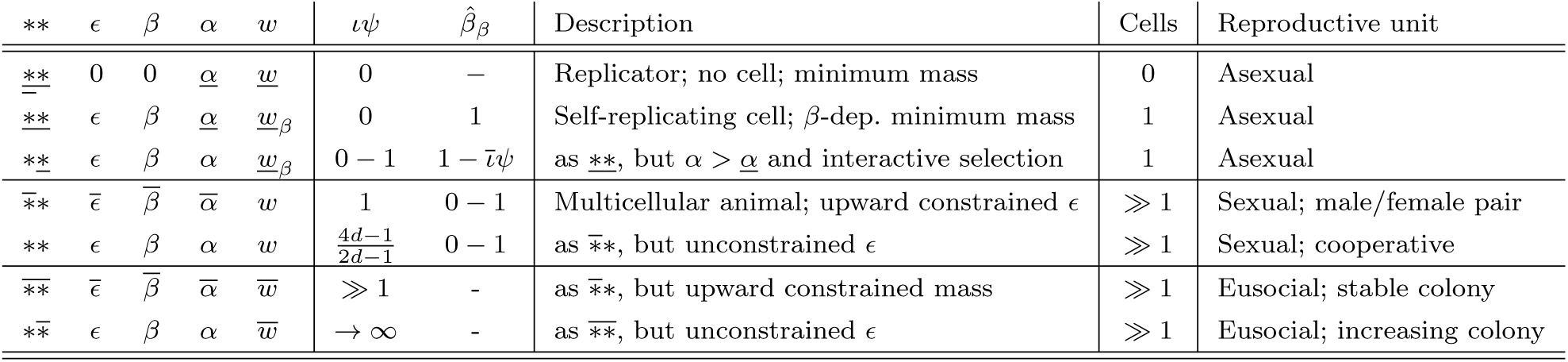
Selection attractors. The asterisk (**) superscripts and main parameters of the selection attractors that evolve from a gradual unfolding of population dynamic feed-back selection from density dependent interactive competition. The bar notation on the left * describes the selection status of *ε*, *β* and *α*, and that of the right * the selection status of *ω*. Underlined asterisks denote a downward minimum or selection constraint, overlined an upward constraint, and unlined no constraint.

The density regulation is partitioned *f* = *f*_*e*_*f*_*ι*_*f*_*s*_ into population exploitation (*f*_*e*_),interference (*f*_*ι*_), and the local exploitation of the individual (*f*_*s*_). The exponents of the mass-rescaling allometries are then following from a selection optimisation of the ecological geometry of the foraging process that defines the three regulation components *f*_*e*_, *f*_*ι*_ and *f*_*s*_ (Witting 1995, 2016a). These frequency independent equations are also used as the starting point to describe the density-frequencydependent bias in the net assimilated energy across the heritable variants in the population [see e.g. eqn 13]; a bias that is selecting for large body masses and major life history transitions (Witting 1997, 2002a).

Irrespectively of the underlying details of the natural selection of mass, the average mass of a selection attractor can be expressed as

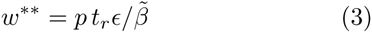

from a rearrangement of eqn 1. And when the theoretically deduced allometric relations of Witting (2016a) are inserted for the parameters on the right hand side of eqn 3, we find the body mass of the evolutionary equilibrium

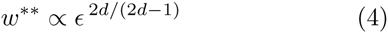

to be expressed as an allometric function of the net assimilated energy. This reects a general principle where natural selection can select for a non-negligible body mass only when the organism has sufficient net energy available for the development of the offspring.

## 3 Replicating molecules

My starting point for evolution by natural selection is the origin of replicating molecules with no intrinsic metabolism (*β* = 0), no cell and practically no mass (<u>*ω*</u>), and a replication that is driven directly by an extrinsic source of energy (metabolism), with the rate of self-replication (eqn 1) being zero as *ε* = 0. While the replication of these molecules requires energy, they are essentially zero-energy replicators as they have no intrinsic metabolism to generate energy.

The molecular replicator may evolve to a self-replicating cell with an internal metabolism that generates net energy for self-replication, with the *β* = 0 ↔ *β* > 0 transition separating life histories with no energy (*ε* = 0) from those with energetic states (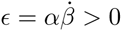).

### 3.1 Initial metabolism depends on mass

The initial selection of metabolism and mass may occur by a dependence of mass specific metabolism on mass when mass is close to an absolute minimum (DeLong et al. 2010). This is because the metabolism that generates the net energy that is required for self-replication is dependent at least upon the mass of the involved metabolic molecules.

Extremely low levels of metabolism are not necessarily dependent upon the development of a compartment like a cell. But it is assumed here that a self-replicator with an advanced form of metabolism is dependent upon a cell where the different molecules of the metabolic pathways can concentrate (e.g., Oparin 1957; Miller and Orgel 1974; Maynard Smith and Szathmáary 1995; Michod 1999; Wächtershäuser 2000; Koch and Silver 2005). Then, with selection driven by the self-replication energy that is generated by metabolism, it follows that the evolution of the self-replicating cell is the evolution of a metabolic compartment. The increase in net energy for self-replication with increased mass specific metabolism can then be the selection that drives the evolution of the cell and all its associated mass; let it be the mass of the cell membrane, over the heritable code, to the metabolic molecules themselves.

The initial evolution of mass specific metabolism is thus linked to the evolution of the cell and its mass, simply because the metabolism cannot evolve without the mass. This mass dependence of mass specific metabolism is not expected to vanish immediately with the evolution of a self-replicating cell, because larger cells allow for the evolution of more fully developed metabolic pathways. The mass dependence, however, will eventually vanish with the evolution of complete metabolic pathways in larger cells.

To describe this in more detail let us assume that the smallest self-replicators have a passive interactive competition that cannot generate anything but an insignificant bias in the distribution of resources over mass, given the level of interference in the population. Body mass selection is then frequency independent, and the selection gradient across self-replicating variants with a given mass specific metabolism (subscript |*β* is negative

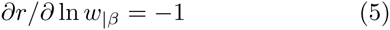

because of the quality-quantity trade-off 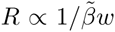 of eqn 1. For this selection, let 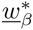 be a partial selection attractor that is defined by the minimum mass (*<u>w</u>*) that is required to sustain a given mass specific metabolism *β*.

A low-energy self-replicator that is evolving a cell with given metabolic pathways may thus be considered to evolve along a distribution of *β*-dependent minimum masses (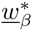), with each mass being selected by eqn 5 as the minimum mass that is required to sustain a self-replicator with the given mass specific metabolism. Smaller masses cannot sustain the metabolism, and larger masses with the same mass specific metabolism are selected towards the *β*-dependent minimum mass because of the quality-quantity trade-off.

As a more developed metabolism requires more mass from the molecules that organise the cell, its metabolism and heritable code, it follows that mass specific metabolism will increase with an increase in the *β*-dependent minimum mass, i.e.,

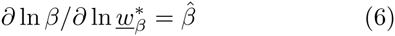

where 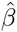 is a positive exponent that declines as the increase in the *β*-dependent minimum mass provides more and more fully developed metabolic pathways.

An initial selection increase in net energy by an increase in metabolic pace will thus be inherently linked to an increase in the *β*-dependent minimum mass. But when is the positive dependence of mass specific metabolism on minimum mass strong enough to generate the extra net energy that is required, not only for the production of more offspring, but also for the production of larger offspring; as required before a selection increase in metabolism and mass can occur?

### 3.2 The initial selection

To examine this selection we have 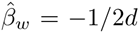 as the mass-rescaling (subscript *ω*) exponent for mass specific metabolism (Witting 2016a). We may then define the dependence of mass specific metabolism on the *β*-dependent minimum mass from the pre-mass relation

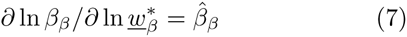

where 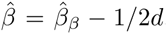, and the pre-mass (subscript *β*) exponent (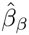) for mass specific metabolism is declining monotonically with an increase in 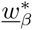, from an initial value (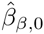) at the lower mass limit (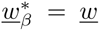) that defines the transition between the replicator with no metabolism and the smallest self-replicator with an initial metabolism.

To transform eqns 5 and 7 into an initial selection gradient for the joint evolution of metabolism and mass, we note from Witting (2016a) that some of the allometric correlations among the different traits cancel each other. This allows for a simpler expression of the selfreplication (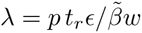, eqn 1) that generates the frequency independent selection on metabolism and mass. With the pre-mass component of survival being proportional to the pre-mass component of mass specific metabolism for all allometric solutions (*p*_*β*_ ∝ *β*_*β*_ ∝ 1/*τ*_*β*_, Witting 2016a), it follows that *pt*_*r*_ reduces to *p*_*ω*_*τ*_*ω*_*τ*_*r*_. Then, given mass invariance for *p*_*ω*_, *τ*_*ω*_, *τ*_*r*_ and 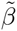 (Witting 2016a) we have *λ* ∝ *τ*_*ω*_*ε*/*ω*, that reduces to *λ* ∝ *β*_*β*_/*ω* given *ε* ∝ *β* ∝ *β*_*β*_*β*_*ω*_, *τ*_*ω*_ = 1/*β*_*ω*_, and 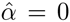 initially. And with *r* = ln *λ* and the pre-mass component of mass specific metabolism being defined by eqn 7, we obtain the following fitness profile

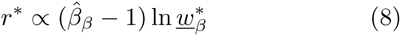

and selection gradient

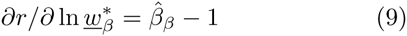

on the *β*-dependent minimum mass.

### 3.3 The evolution of virus

Before we use eqn 9 to predict the evolution of self-replicating cells with internal metabolism, let us consider the other side of the coin that is an evolutionary dead-end that maintains molecular replicators with no metabolism as the selection attractor at the cost of the evolution of self-replication, metabolism and mass.

For a virus-like replicator with no internal metabolism to persist as a selection attractor it is required that the self-replication selection at the potential transition between the molecular replicator with no metabolism and the smallest self-replicator is selecting for the molecular replicator at the cost of a self-replicator with an internal metabolism. This selection occurs whenever the maximum exponent 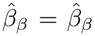 of eqn 9 is smaller than one. The selection gradient is then negative with constant selection for a joint decline in mass and mass specific metabolism.

Unlike selected self-replicators, these replicators cannot increase replication by an increase in mass specific metabolism. This is because the energetic demands for the mass of the offspring increases linearly with mass, while the metabolism that generates the required energy can only increase sub-linearly with mass. The replicators may instead increase their replication rate by a mass that evolves towards zero, with the side effect that they are shutting down their metabolic processes. The final attractor

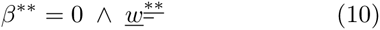

is the molecular replicator with no internal metabolism, no cell, practically no mass, no self-replication, and a replication that is driven by an extrinsic source of metabolism [for 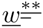, the bar under the last asterisk denotes the negative selection of eqn 5 on *ω*, and the double bar under the first asterisk denotes the lower bound on *ε* with no metabolism (*β* = 0) and a passive resource handling (*<u>α</u>*)].

Molecular replicators with no metabolism and practically no mass may thus not only be the potential ancestor for more complex self-replicators with an internal metabolism, but they may also be selection attractors of self-replication selection itself.

Although extremely small, viruses are likely significantly larger than the replicating molecules that were formed prior to the evolution of self-replicating cells with an internal metabolism. These relatively large masses of viruses are expected to be selected by an energy-assisted-replication that is fuelled by the metabolism of their host. Because of this increased mass, the maximum metabolic exponent (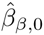) may be smaller in viruses than in an initial molecular replicator that was formed by other processes than replication. It is thus quite likely that viruses have masses that are situated at a joint attractor, where they are selected not only by energy-assisted-replication, but also by the self-replication selection that occurs in viruses that mutate to some initial forms of intrinsic metabolism. Having no metabolism and a time-scale that is defined by the metabolism of their host, viruses are selected beyond the allometric model in Witting (2016a).

## 4 Minimum self-replicating cells

Let us now consider what happens when the maximum pre-mass exponent for mass specific metabolism (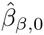) is larger than one.

### 4.1 The initial metabolism and mass

Given replicating molecules that are smaller than viruses, it is likely that at least some of these will have a maximum pre-mass exponent that is larger than one (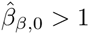). The only thing that is required is the potential for heritable mutations towards metabolic pathways of self-replication where the attachment of catalysts to an initial self-replicator is causing an increase in mass specific metabolism that is about proportional to the increase in mass [Recall that the 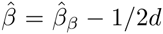 exponent that defines the final relationship between *β* and *ω* is smaller than the 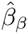 exponent in eqn 9]. When this is the case we find from eqns 7 and 9 that the selection attractor is an equilibrium *β*-dependent minimum mass 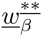 that is defined by a pre-mass exponent of unity

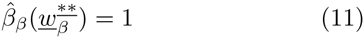

with the single bar under the *ε* (first) asterisk denoting minimum handling (*<u>α</u>*) with internal metabolism (*β* > 0). As this is an equilibrium of frequency independent selection we expect a straightforward relationship between the fitness profile and the selection integral, as illustrated by Figs. 1a, b and c.

**Figure 1:**
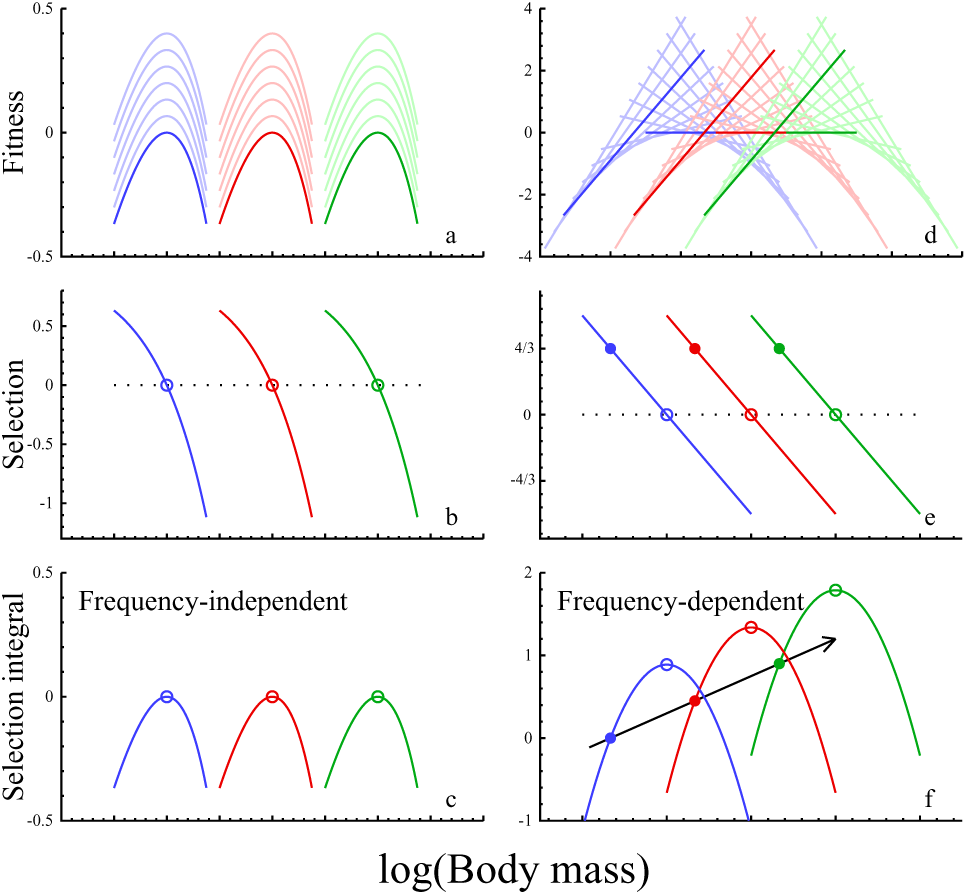
Body mass selection. Fitness profiles/landscapes [**a** & **d**; 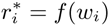, intra-population variation(subscript *i*) in populations that are in dynamic equilibrium], selection gradients [**b** & **e**; 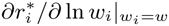 across population variation (no subscript)] and selection integrals [**c & f**; 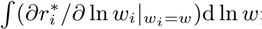 across population variation] for the frequency independent selection of the physiology (**a, b & c**; calculated from model in Section A.1) and the frequency dependent selection of density dependent interactive competition (**d, e & f**; from eqns 21 to 23, given 2D interactions). Selection integrals look like fitness landscapes for physiological selection, but they cannot be visualized from the fitness landscapes of density dependent interactive competition. Equilibrium attractors are shown by open circles (different colours different attractors), and unconstrained selection by interactive competition has steady state attractors (solid circles) with selection integral evolution (black arrow) and exponential increase in net energy and mass (Witting, 2016b). The multiple fitness profiles per colour represent populations with different average variants, with clear coloured curves having average variants at evolutionary equilibrium or steady state.

We may thus conclude that when the maximum pre-mass exponent for mass specific metabolism at the lower mass limit is larger than one (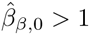), it follows that the net energy that is obtained by the increase in mass specific metabolism increases stronger than linearly with mass, and this will generate an increase in the self-replication rate with mass. This selection for an increase in metabolism and mass will continue until the pre-mass exponent has declined to unity, and the net energy that is generated from extra metabolism is exactly balancing the selection of the quality-quantity trade-off (eqn 5).

### 4.2 Life history

In relation to the life history of these minimum sized self-replicators, we note that for cases where the maximum 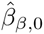 exponent is only slightly larger than one, we might expect the evolution of self-replicators with simple metabolic pathways and no cell. But for larger 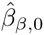 exponents we can expect selection for the formation of a cell-like structure with an increased mass and an internal metabolism.

As it is the formation of the cell that is structuring the metabolic pathways, and as these self-replicators are selected to have the minimum possible mass that is required for their metabolism, they are selected to have a single cell only. And with a metabolic pre-mass exponent (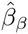) of unity, the selection attractors are so small that their metabolic pathways are incomplete in the sense that it is biochemically possible to increase the metabolic efficiency per unit mass.

A 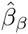 exponent of unity implies also that the body mass exponent for resource handling (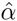) is predicted to be zero (Witting 2016a). This indicates that these self-replicating cells have a passive handling of their resources. And as the selection attractors will have no significant resource bias from interactive competition, they have frequency independent selection for asexual reproduction (Witting 1997, 2002a).

### 4.3 Allometries

With the dependence of mass specific metabolism on mass being different for different metabolic pathways, and the dependence evolving with the evolving self-replicator, we can expect low-energy self-replicators with different minimum masses because of adaptations to a variety of niches and metabolic pathways. And with these attractors being predicted as the smallest group of self-replicating cells, they would resemble prokaryotes. The predicted allometric exponents are given by the 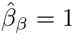 rows in Table 2 in Witting (2016a). For behaviour in three spatial dimensions, this implies a mass specific metabolism and a population dynamic growth rate that increase to the 5/6 power of mass.

Empirical exponents in heterotrophic prokaryotes (DeLong et al. 2010) around 0.84 (0.96±0.18 for active; 0.72±0.07 for inactive) for mass specific metabolism, and 0.73 for the rate of population growth, suggest that the body masses of prokaryotes may indeed evolve from a positive dependence of mass specific metabolism on the mass of the self-replicating cell.

## 5 Interacting self-replicating cells

Minimum sized self-replicators that are larger than the attractor of eqn 11 will have an increased mass specific metabolism and more net energy available. But they cannot be selected exclusively by the dependence of mass specific metabolism on mass.

### 5.1 Evolution of interactive competition

As there seems to be no physiological constraint that will generate frequency independent selection for a general increase in mass (Appendix A), let us examine how the emergence of a resource bias from interactive competition is affecting the selection of metabolism and mass.

This frequency dependent selection reects that the larger individuals in a population may dominate the smaller individuals during competitive encounters, at least when other things are equal. This is expected even for the passive type of behaviour that we may expect in simple self-replicators, where larger individuals will have more kinetic energy than smaller individuals and thus also a higher probability of winning a competitive encounter. This will create a bias (Δ*μ*_*i*_ = *μ*_*i*_ − *μ*) in the fitness cost (*μ*) of a competitive encounter, where Δ*μ*_i_ is the cost of the *i*th variant relative to the average cost (*μ*) in the population. With a symmetrical cost bias on logarithmic scale, and *ψ* being the gradient that measures the fitness cost per unit interference on log scale across the body mass variation in the population, we find that the differentiation in the cost of interference per unit interference is

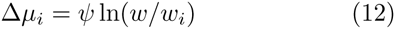

Here it is the mass (*ω*^*i*^) of an individual (*i*) relative (*ω*_*i*_/*ω*) to the average mass (*ω*) that defines the ecological dominance of the individual in the population, and this translates into the fitness profile

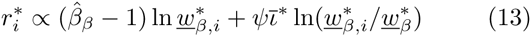

for the potential evolution of minimum masses 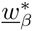, with the cost of interaction increasing with a level of intraspecific interference (*ι*) that is maintained at its maximum [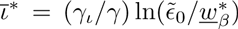 from eqn 17 in Witting 2016a] by the *β*-dependent minimum mass.

The selection gradient on the cost gradient

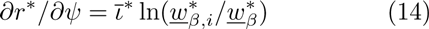

of eqn 13 is positively related to the level of interference, with disruptive selection for a larger *ψ* in the individuals that are larger than the average, and a smaller *ψ* in the individuals that are smaller than the average. This reects a selection battle, where the larger than average individuals are selected to monopolize the resource during encounters, and the smaller than average individuals are selected to avoid the monopolization of the larger individuals. While the avoidance of smaller individuals may limit the average gradient to some degree, the energetic dominance of the larger individuals sets a natural upper limit to this avoidance. It is thus the monopolization component that will dominate the evolution of the cost gradient, with mass being selected as a trait that is used to dominate other individuals during interactive encounters.

### 5.2 Initial interactive selection of mass

To describe this interactive selection of mass, from the fitness profile of eqn 13 we obtain the selection gradient across the intra-specific variation in the *β*-dependent minimum mass

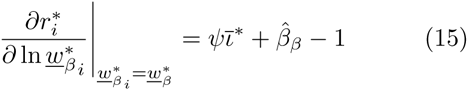

The evolution of an interacting self-replicator with a pre-mass exponent that is smaller than one is thus dependent upon the evolution of a maximum resource bias that is larger than zero (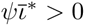). The evolution of such a gradient will generate a selection attractor with a *β*-dependent minimum mass that is defined by a pre-mass exponent for mass specific metabolism that is smaller than unity

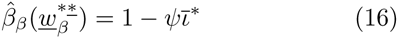

This evolution can continue along a gradient where the maximum cost bias increases towards unity, reecting a population dynamic feed-back selection that is unfolding from an increase in the net energy of the average individual in the population. The feed-back, however, is not yet fully developed, and body mass selection is still dependent upon the frequency independent dependence of mass specific metabolism on minimum mass.

### 5.3 Life history

From eqns 7, 11, and 16 we find that the interacting self-replicators are larger than self-replicators with no interactions, and that they increase in size with an evolutionary increase in the maximum resource bias (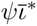) of interactive competition. As prokaryotes, they are selected to have the minimum possible mass that is required for their metabolic functioning, and they are thus selected to be unicellular self-replicators. And being predicted as a second size class of unicellular attractor beyond replicating molecules, we expect the interacting self-replicating cells to resemble larger unicells like protists and protozoa.

We note also that the frequency dependent selection from the unfolding feed-back is dependent on net energy that is generated from an independent selection on resource handling. This is because of the 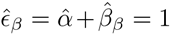 relationship from eqn 29 in Witting (2016a). This implies that the evolution is dependent upon the evolution of a proportional relationship between mass and the pre-mass component of net energy. And with the 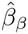 exponent being smaller than one, it follows that the exponent for resource handling (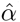) is selected to be larger than zero.

It thus seems that an evolutionary increase in resource handling is needed to maintain the frequency dependent selection of interactive competition that is required for an evolutionary increase in the mass of the interacting self-replicating cell. Hence, the evolution of protozoa-like cells seems to require both increased resource handling and an interactive competition that generates some bias in the resource distribution over mass. Protozoa and other larger unicells are thus predicted to have a more organized and developed resource handling and interactive behaviour than the individuals of prokaryotes.

As these larger unicellular self-replicators are predicted to have a maximum resource bias that is smaller than unity (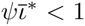), it follows that their selection for major life history transitions is dominated by the frequency independent selection of their physiology (Witting 1997, 2002a, 2008). This implies that they are selected to have asexual reproduction. But as the transition boundary for the frequency dependent selection of multicellular organisms with sexual reproduction lies at a maximum resource bias of unity (Witting 1997, 2002a, 2008 and next section), we may also expect that some of the larger interacting self-replicators with the strongest resource bias can be selected to show some initial traces of both multicellularity and sexual reproduction.

### 5.4 Allometries

To examine the selection of allometries, recall that the variation in body mass across populations of interacting self-replicating cells are predicted to evolve from the coevolutionary continuum in net energy and behavioural interactions that will make the maximum cost bias (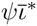) increase from zero to one. Hence, from eqn 16, we predict a joint evolutionary increase in mass and the pre-mass component of mass specific metabolism, with the pre-mass exponent for mass specific metabolism (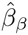) declining from one to zero. So instead of having a continued increase in mass specific metabolism with mass, as predicted for the minimum self-replicator, we find from Table 2 in Witting (2016a) that interacting self-replicators are predicted to have an initial increase in mass specific metabolism with mass (5/6 exponent in 3D, and 3/4 exponent in 2D), that will decline with mass first to a mass invariance, and then to a negative correlation with an exponent that approaches −1/6 in 3D, and −1/4 in 2D, as the dependence of mass specific metabolism on mass is vanishing with the evolutionary increase in mass.

By excluding the four smallest protozoa with exceptionally high metabolic rates from the data of Makarieva et al. (2008), and by least-squares fitting a third degree polynomial to the remaining data for inactive protozoa (*n* = 48), Witting (2016a) obtained point estimates of the body mass exponent for mass specific metabolism that declined from 0:61 across the smallest [log *ω*(kg) = −13.5], over zero across intermediate [log *ω*(kg) = −11], to a minimum of −0.20 among the largest protozoa [log *ω*(kg) = −8.0, see Fig. 6]. This change indicates that protozoa may be selected by an interactive feed-back selection that is unfolding with an increase in the maximum cost bias.

## 6 Multicellular animals

A next transition in the evolution of metabolism and mass occurs when the maximum cost bias of interference competition evolves beyond unity (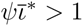). This will cause the unfolding of a fully developed population-dynamic feed-back selection that selects for organisms that are larger than the minimum that is required for their metabolic functioning. This makes the 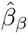 exponent functionally independent of mass, and it allows for an in principle independence between the evolution of mass, resource handling, and mass specific metabolism. The three traits will however be interlinked by mass-rescaling and population dynamic feedback selection, where the selection of mass is dependent on the net energy that is generated by resource handling and metabolic pace.

### 6.1 Theoretical background

The idea that interactive competition may be the main cause for the selection for large body masses was first considered with density independent interference. This can generate evolutionary arms races with frequency dependent selection for a directional evolution where an interactive trait, like body mass, continues to absorb energy from the reproductive rate (Simpson 1953; Dawkins and Krebs 1979; Parker 1979; Haigh and Rose 1980; Maynard Smith 1982; Parker 1983; Brown and Maurer 1986; Maynard Smith and Brown 1986; Vermeij 1987; Härdling 1999). The intrinsic population dynamic growth rate and the population dynamic equilibrium will then decline towards zero, with the population evolving towards extinction.

The directional evolution of arms races applies when the level of interactive competition is sufficiently large and density independent, but selection from intraspecific interactive competition is density dependent, and this needs to be incorporated explicitly into the model of natural selection (e.g. Abrams and Matsuda 1994; Witting 1997; Heino et al. 1998; Kisdi 1999; Gyllenberg and Parvinen 2001; Dercole et al. 2002; Dieckmann and Metz 2006; Rankin 2007; Matessi and Schneider 2009; Peischl and Schneider 2010). This density dependence implies that the relative fitness of a variant is density-frequency-dependent, with the cost of interference depending not only on your own phenotype, but also on the phenotypes of your opponents, and on the number of interactive encounters that increases monotonically with density. And this generates feed-back between the life history and the population dynamic selection pressure on the life history.

### 6.2 Population dynamic feed-back selection

The fully developed population dynamic feed-back selection is illustrated in Fig. 2. It emerges from the pre-mass selection on the resource handling (*α*) and metabolic pace (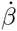) components of net energy (*ε*). This defines the growth (*r*) potential of the population, with density regulation (*γ*) determining the equilibrium abundance (*n*\*) and the level of interactive competition (*ι*\*) that generates the selection pressure on mass (*ω*). The inner loop of the figure is the mass-rescaling that is described by Witting (2016a), where metabolic trade-off selection is adjusting the physiological components of metabolism and time (*t*_*j*_, *β* & *t*_*r*_) in response to the selection on mass, and the optimisation of density regulation is adjusting the home range (*h*) to the changes in mass, metabolism and time.

**Figure 2:**
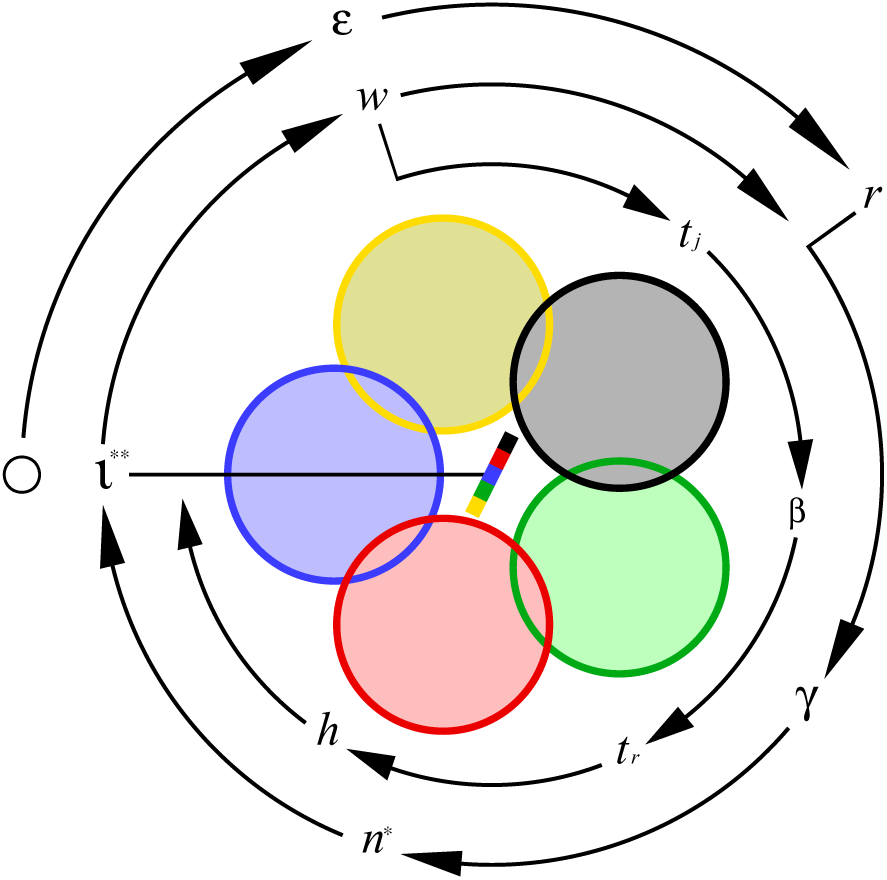
An illustration of population dynamic feed-back selection. Symbols relate to the population average, and coloured circles symbolize individual home ranges in two-dimensional space with interactive competition in zones of overlap. Winners (dominating colour) monopolize resources, and this generates a body mass biased resource access that is proportional to the slope of the multi-coloured bar in Centrum, with the invariant interference (*ι*^**^) of the selection attractor determining the evolution of this bias. The attractor is given by the feed-back selection of the outer ring of symbols [*r*:population growth → *γ*:density regulation → *n*^*^:population abundance → *ι*:interference level → *ω*:selection on body mass → *r*:population growth]. Selection for a change in mass initiates the inner loop of massrescaling selection [*ω*:mass change → *t*_*j*_:juvenile period → *β*:metabolic rate → *t*_*r*_:reproductive period → *h*:home range → *ι*:interference], with both loops adjusting to the invariant interference attractor (see Fig. 3). The black o to the left represents the origin of self-replication, and selection for an exponential increase in net energy (*ε*) maintains a relatively high *r* and continued feed-back selection for an exponential increase in mass.

### 6.3 Competitive interaction fix-points

With a body mass that is selected beyond the minimum that is required for a given mass specific metabolism, the intrinsic dependence of mass specific metabolism on mass is vanishing, and the fitness profile on body mass reduces to

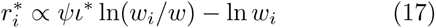

with the following selection gradient

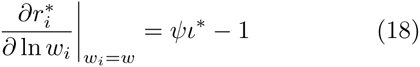

The equilibrium attractor of population dynamic feedback selection is thus a competitive interaction fix-point

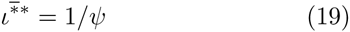

with an invariant level of interference competition as the population condition for a neutrally stable selection the population condition for a neutrally stable selection is exactly so high that the increased reproductive potentials of the smaller individuals are balanced against the resource monopolization of the larger individuals so that fitness is similar across variation in mass.

The 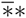-attractor of eqn 19 is the result of a stable net energy 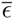 that is upward constrained for some reason. The **-attractor for unconstrained selection with an exponential increase in *ε* is characterized by a somewhat higher level of invariant interference

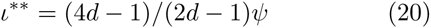

and an exponentially increasing mass (Witting 1997, 2003).

The competitive interactions fix-points of eqns 19 and 20 are based on an unconstrained body mass, where the attracting mass (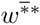 & *ω*^**^) is an energetic buffer that absorbs the energy that is initially selected into enhanced reproduction and population growth. If the mass evolves to an upper bound (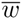) it follows that this energy can no longer be absorbed by selection for mass, with the result that the population density and the level of interference competition evolves to a much higher stable (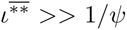) or increasing (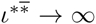) level, dependent upon a stable or increasing net energy (Witting 1997, 2003).

How may these competitive interaction fix-points coevolve with the selection of optimal density regulation (Witting 2016a), that is adjusting the home range to optimise the foraging behaviour as a balance between the cost of interference competition and the cost of the local resource exploitation of the individual. As this density regulation optimum depends mainly on behavioural traits it should respond relatively fast to natural selection, and let us therefore consider a population at the regulation optimum. If the level of interference at this optimum is higher than expected from the interaction fix-point, we have interactive selection for an allocation of energy from replication to mass, with a decline in abundance and interference towards the interaction fix-point. Now, as the density regulation optimum is invariant of the abundance (eqn 18 in Witting 2016a), we find the optimal home range to be unaffected by this selection. The selection decline in interference is then inducing a decline in interference regulation (increase in *f*_*ι*_) that alters the absolute value of the joint regulation (*f*_*ι*_*f*_*s*_) with local resource exploitation (*f*_*s*_), but not the optimal home range (Fig. 3).

**Figure 3:**
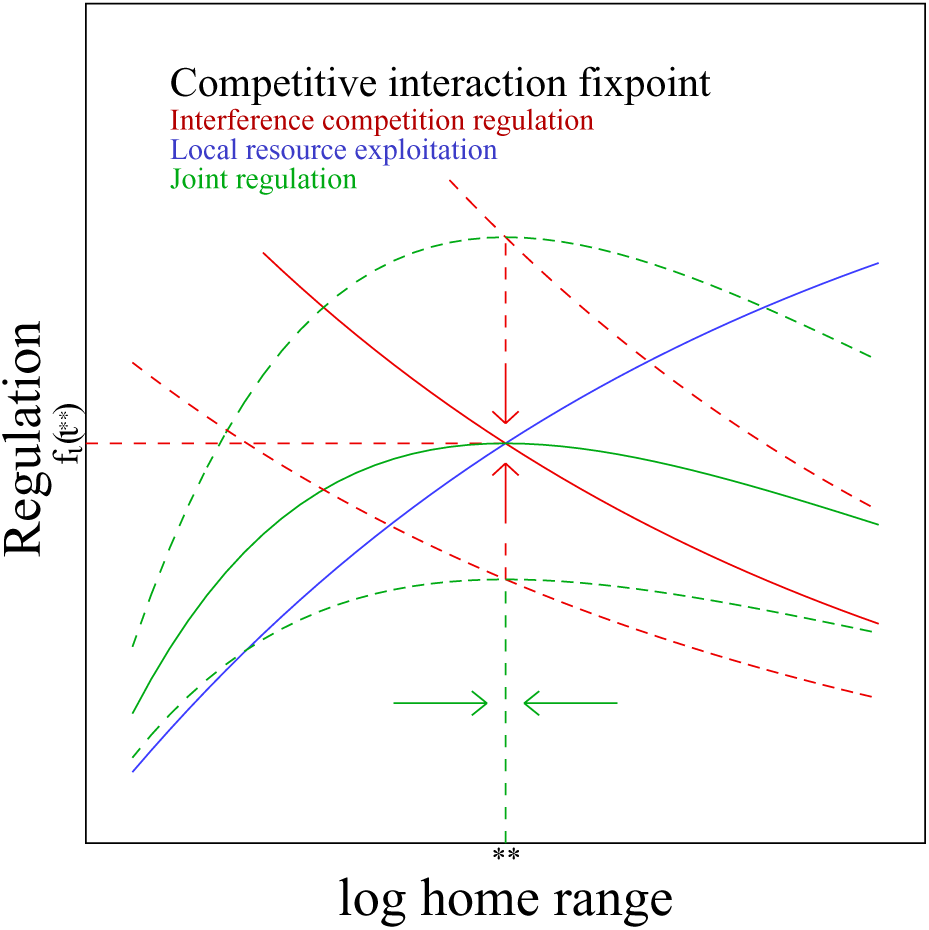
The density regulation attractor. The home range of optimal density regulation (**), as defined by the joint regulation (*f*_*ι*_*f*_*s*_; green curves) of interactive competition (*f*_*ι*_; red curves) and local resource exploitation (*f*_*s*_; blue curve). The optimal home range (eqn 18 in Witting, 2016a) is independent of the feed-back between the population abundance and the interactive selection on mass. And with the level of interference being dependent on abundance (eqn 16 in Witting, 2016a), it follows that the density regulation of interference competition is adjusted by body mass selection to a joint selection attractor (solid red and green curves), where regulation at the home range optimum coincides with the regulation [*f*_*ι*_(*ι*^**^) of the competitive interaction fix-point (*ι*^**^) for the selection attractor on mass.

The density regulation of interference [*f*_ι_(ι)] is thus adjusted by the population dynamic feed-back selection to a joint selection optimum, where the interference regulation of the home range optimum coincides with the regulation [*f*_ι_(ι^**^)] of the competitive interaction fixpoint (e.g., *ι*^**^).

### 6.4 Interactive selection of mass

It is possible to visualize the mass attractor of population dynamic feed-back selection by including the population dynamic feed-back of density dependent interference competition (eqns 17 and 80 in Witting 2016a) into the equations. The fitness profile (Fig. 1d) on body mass is then

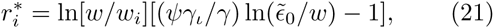

and the selection gradient (Fig. 1e)

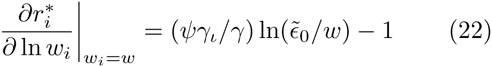

with 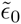 being a measure of net energy on the per generation time-scale. The latter equation is then integrated across the potential evolution of the average variant in the population to obtain the selection integral

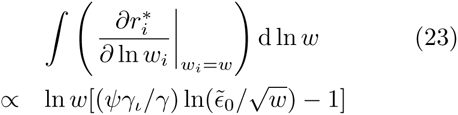

that illustrates the evolutionary attractor (Fig. 1f). This selection by density dependent interactive competition has no absolute measure of relative fitness, and no concept of an increase in average fitness. The result is a fitness landscape (profile) that has no direct visualization with the attractor of the selection integral (compare Fig. 1d & 1f).

The attracting mass (eqn 3) implies an equilibrium mass that reects energy that is initially allocated into enhanced population growth and thereafter selected into increased mass by interactive competition, with the resulting level of interference competition being invariant of mass.

### 6.5 Life history

The transition in the interactive competition to a maximum cost gradient above unit is not only selecting for a transition from self-replicators with the minimum masses that are required for their metabolic functioning, to larger organisms where energy is selected into mass to enhance the competitive quality of the individual. With a mass that is no longer selected to the minimum that is required to sustain metabolism, there is no longer direct selection for the existence of only a single metabolic compartment (one cell).

In the absence of direct selection for single-celled individuals, co-operation at a higher multicellular level may trade-off fitness at the cellular level to enhance the overall fitness of the organism (Buss 1987; Michod 1996, 1997, 1999; Michod and Roze 2001). The increased functionality of resource handling, interactive behaviour and resource transportation that can be obtained from a division of a single large cell into a multitude of cooperating cells with smaller masses, is then expected to select for the emergence of multicellular organisms.

The selection of other life history traits is also dependent on the level of interference in the population, with the transitions between the different competitive interaction fix-points selecting for major life history transitions (Witting 1997, 2002a, 2008). The increase in the maximum cost gradient above unity is likely to induce selection for a soma with senescence, where it is beneficial to take energy from self-repair and use it in interactive competition (Witting 1997, 2008). And it is also selecting for the evolution of sexual reproduction between a male and female individual, where the male is specialising in interactive competition at the cost of self-replication, and the female is sharing her genome in the offspring with the male in order to attract the competitively superior males (Witting 1997, 2002a, 2008). The result is interactive competition that is generating frequency dependent selection for the evolution of sexual reproduction with a diploid genome.

This selection for a reproducing unit with specialised multicellular individuals and sexual reproduction is unfolding to higher levels of organisation with an increase in the interactive competition in the population (Witting 2002a). Where the interactive competition of the energetically constrained mass attractor (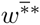) is selecting for pair-wise sexual reproduction, the increased interference at the evolutionary steady state (*ω*^**^) is selecting for cooperative reproduction where a single or a few offspring workers are enhancing the interactive quality of the sexually reproducing pair. And the potentially much higher interference at the upward constrained mass attractors (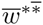 or 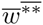) selects for an eusocial colony with thousands of interacting offspring workers; with the energetically constrained attractor (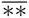) having a stable colony size, while the increasing energy of the energetically unconstrained attractor (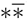) is selected into a continued increase in colony size (Witting 2002a).

Owing to a diminishing return in the interactive quality that the reproducing unit can gain from sexual reproduction between a female and an increasing number of males, these co-operatively and eusocially reproducing units have active selection for pairwise sexual reproduction and offspring workers at the cost of higher levels of sexual reproduction with more than two individuals involved (Witting 2002a). And where the pair-bond in eusocial termite-like species selects for a diploid genome, the absence of a pair-bond in eusocial hymenoptera-like species is selecting for a male-haploidfemale-diploid genome (Witting 1997, 2007).

### 6.6 Allometries

With masses that are larger than the minimum that is required for fully developed metabolic pathways, and the primary selection on metabolism and resource handling being independent of the selection of mass, the allometric exponents for high-energy organisms may in principle take any of the values in Table 2 in Witting (2016a). What is essential for the evolution of the exponents is the underlying causes for the evolutionary variation in the mass of the species that are being compared in the allometric correlation.

If the primary cause for the mass variation is selection on metabolic pace, we expect a pre-mass exponent for mass specific metabolism around one, and a mass specific metabolism that increases to the 5/6 power of mass in 3D, and the 3/4 power in 2D. If on the other hand, we are examining a taxonomic group where species have diversified and adapted to a broad spectrum of niches, we expect that the primary causes for the variation in mass is variation in *α*, i.e., variation in the handling and densities of the underlying resources. The pre-mass and densities of the underlying resources. The pre-mass zero, and we have Kleiber scaling with a −1/4 exponent for mass specific metabolism for two dimensional interactions, and a corresponding exponent of −1/6 across species that forage and interact in three spatial dimensions.

Table 1 lists all the selection attractors on mass together with a brief description of the predicted life histories; ranging from replicating molecules to the eusocial colony of the multicellular animal. The initial conditions that determine the specific attractor for an evolutionary lineage at a given point in time are the selection and constraint status of mass specific metabolism (*β*), resource handling (*α*) and mass (*ω*), and the associated resource bias in the population (*ιψ*).

## 7 Macro evolution

So far we have predicted a macro evolutionary succession towards larger lifeforms, starting from replicating molecules, over minimum self-replicating cells, and larger unicells, to the multicellular animal. We found the progression towards larger size to be driven by primary selection for increased mass specific metabolism in prokaryotes and the smaller protozoa, with the importance of metabolism for the evolution of mass declining with an increase in protozoa mass.

With typical exponents around −1/4 and −1/6 for mass specific metabolism in the major taxonomic groups of multicellular animals, the importance of metabolism for the evolution of mass seems to have vanished when we deal with the evolution of the body mass variation in these taxa. Their body mass distributions seem instead to be dominated by the evolution of divers handling of resources across ecological niches. Yet, selected variation in mass specific metabolism should also generate body mass variation in multicellular animals, and we may therefore ask whether the evolution of metabolism and mass is interlinked in a macro evolutionary pattern with different allometric exponents within and across the major taxa of multicellular animals?

### 7.1 The three mass scaling components

In order to examine this with the proposed model, consider first the predicted relationships between the three mass scaling components (pre-mass, mass-rescaling,and post-mass) for the four cases of pre-mass selection on metabolism that is given by Witting (2016a). This is shown for mass specific metabolism in Fig. 4, with the blue lines being pre-mass selection on metabolism, green being the local rescaling with mass for a given mass, and red being the final post-mass allometry. The pre-mass selection on metabolism is generating the metabolic span of the blue line (in time or across species), and selection on mass is generating the span in mass, with the green lines illustrating local mass-rescaling and the final post-mass mass-metabolism relationship evolving along the red line.

**Figure 4:**
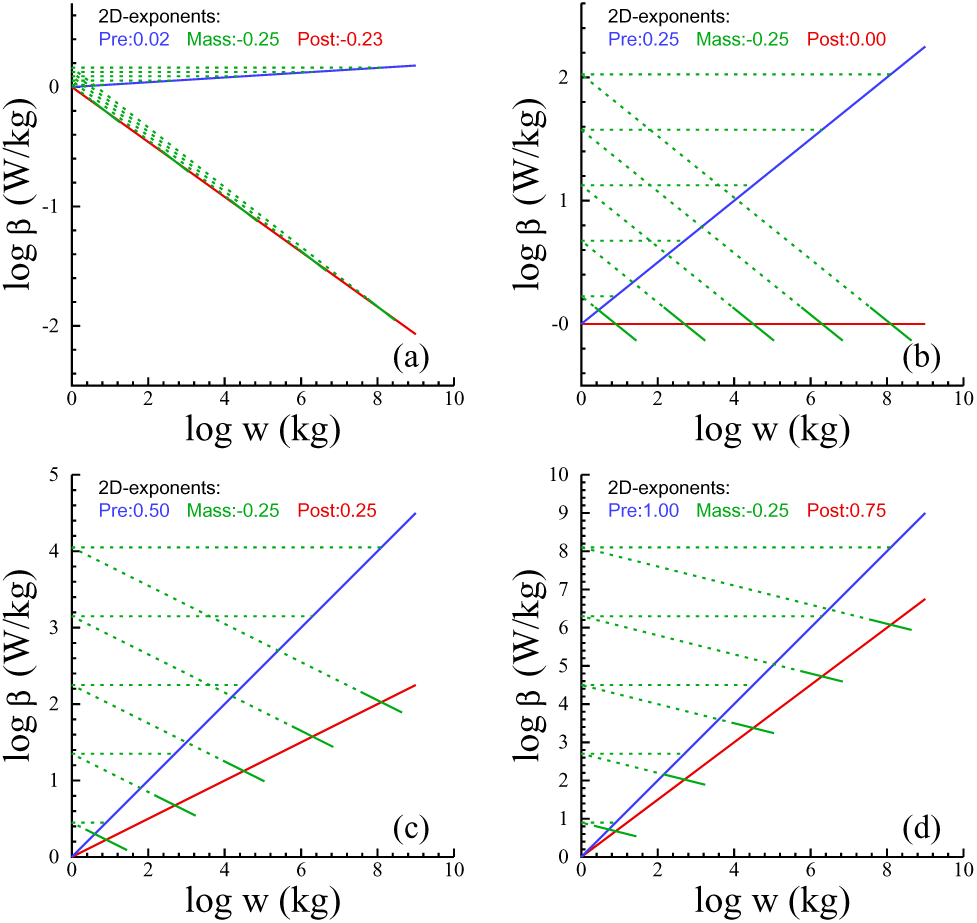
Pre-mass, mass & post-mass scaling. The post-mass allometries for mass specific metabolism (red) evolve from pre-mass selection on metabolic pace (blue) and mass-rescaling (green) from the associated evolutionary changes in mass, given here in plots **a** to **d** as a function of the metabolic-rescaling exponent 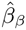 for pre-mass selection on metabolic pace. The dotted green lines illustrate that the metabolic scaling from pre-mass selection (blue) is the scaling of the intercepts of local mass-rescaling (solid green) on the final post-mass allometry (red). Plot **d** represents the limit case where all, and plot **a** where almost none, of the evolutionary variation in mass is induced by the pre-mass evolution of the metabolic intercept (see text for details).

For the 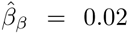 case of Fig. 4a, there is almost no pre-mass evolution in metabolism. The result is a body mass that reects independent selection on resource handling and availability with a post-mass allometry (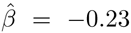) that scales almost as massrescaling (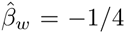). At the other extreme we have Fig. 4d, where 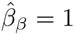 and mass is evolving exclusively in response to the metabolic increase of the blue line, with a post-mass allometry that is positively related to mass 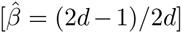 despite of the negative rescaling with mass (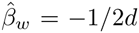). The intermediate 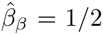 case is shown in Fig 4c, and Fig 4b illustrates 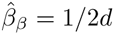 where the evolutionary decline in metabolism caused by mass-rescaling is outbalancing selection on metabolic pace with the final mass specific metabolism being invariant of the evolving mass.

### 7.2 The macro evolutionary pattern

With the typical post-mass scaling of mass specific metabolism in the major taxa of large bodied organisms resembling mass-rescaling with an exponent around −1/2*d*, the local mass-rescaling of the solid green lines in Fig. 4 can be seen as species distributions of different taxa, with the body mass variation within the taxa originating from an evolutionary diversification in the handling of resources across a variety of ecological niches. The red lines of the different plots will then represent different routes for an evolutionary divergence between the major taxa, with the possible macro evolutionary routes between taxa being dependent upon the mode of natural selection on metabolic pace.

This interpretation is illustrated in Fig. 5 together with the predicted positive and intermediate scaling of metabolism with mass in the two lifeforms of self-replicating cells. Included is also the transition from the self-replicating cell with asexual reproduction to the multicellular animal with sexual reproduction as predicted by selection by density dependent interactive competition (Witting 1997, 2002a, 2008). This transition is predicted to lie at the lower boundary where interactive competition can select for a larger than minimum sized organism, which is also the lower boundary of the within taxa −1/2*d*-power scaling of mass specific metabolism with mass.

**Figure 5:**
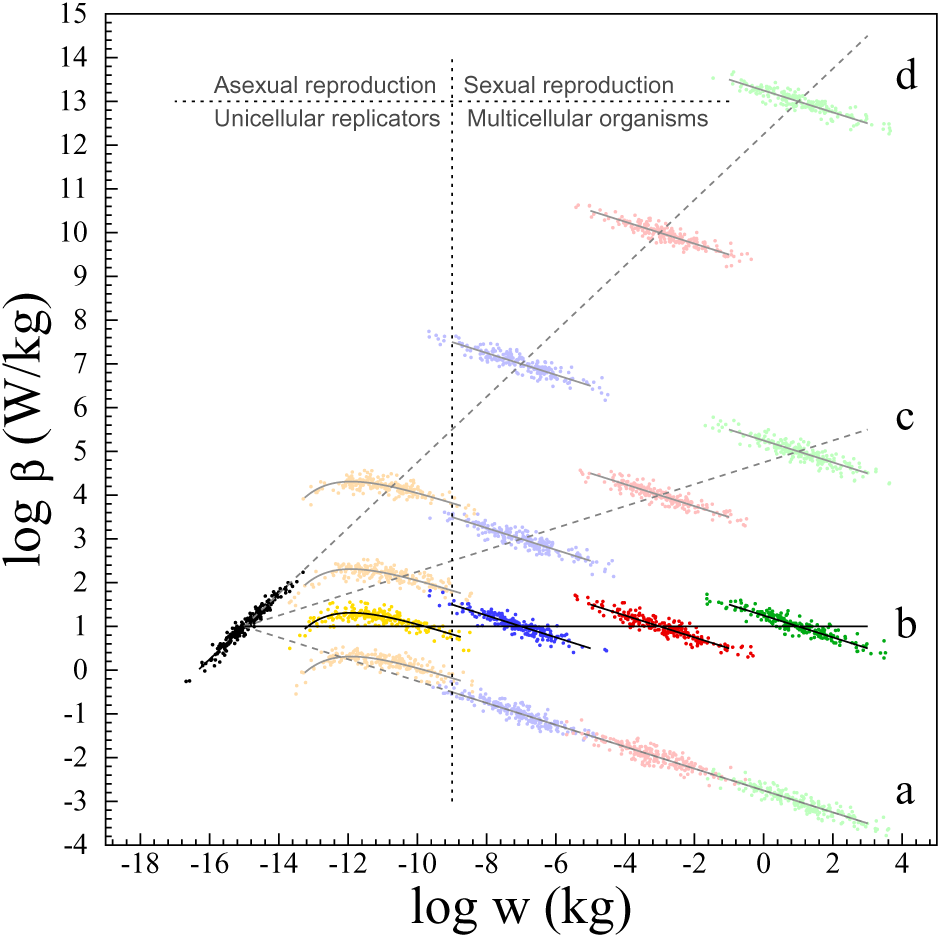
Macro evolution. Theoretical 2D examples for the evolution of major taxonomic groups (different colours), when *i*) prokaryote-like self-replicators (black) evolve by a dependence of mass specific metabolism on mass with 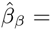 = 1, *ii*) protozoa-like self-replicators (yellow) evolve by the same dependence and interactive competition (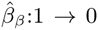), and *iii*) multicellular animals evolve by interactive competition with evolutionary diversification across ecological niches (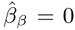). Macro evolution across taxa follows **a**) if there is no pre-mass evolution in 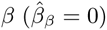, **b**) if *β* evolves along an upper bound (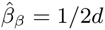), **c**) if metabolic pace and resource handling are equally important for the evolution of net energy (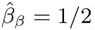), and **d**) if there is no pre-mass evolution in resource handling (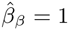). The predicted transition from the asexual self-replicating cell to the multicellular animal with sexual reproduction is also illustrated (for details see the text).

If the 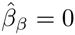 condition that explains the typical postmass scaling within the taxonomic groups of larger animals were to apply also across taxonomic groups, we should see a single 1/2*d*-like scaling of mass specific metabolism across all larger species (Fig. 5a). But a global 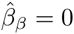 exponent is not our first expectation because, contrary to the prediction of eqn 20 in Witting (2016a), this would imply a general absence of natural selection on metabolic pace. A 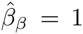 exponent across the major taxa (Fig. 5d) is also unlikely as it would imply that all the variation in mass between the major taxa would be induced by selection differences in metabolism, while all the mass variation within the different taxa of large bodied organisms would be induced by selection differences in resource handling and availability. A more likely base-case could be 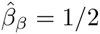 (Fig. 5c), where natural selection on pace and handling is almost equally important at the scale of the major taxa.

But maybe most likely at the macro evolutionary scale is 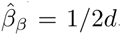. This scenario is expected because we have already predicted that the metabolic pathways should be fully developed at the evolutionary transition to the multicellular animal. Macro evolution across taxa of multicellular animals may then proceed along an upper bound on mass specific metabolism, where an evolutionary increase in mass would require a primary increase in resource handling and/or resource availability. This would generate a mass-rescaling with an allometric downscaling of mass specific metabolism and, if handling increases sufficiently slowly, we should expect a subsequent increase in mass specific metabolism due to pre-mass selection on biotic pace. The overall macro evolution of mass specific metabolism would then proceed along a metabolic bound as in Fig. 5b, while evolutionary diversification across niches would generate −1/2*d*-power scaling within large bodied taxa that expand in species numbers across ecological niches.

Patterns of mass specific metabolism within and across major taxa from prokaryotes to mammals have been studied by Makarieva et al. (2005, 2008), DeLong et al. (2010) and Kiørboe and Hirst (2014), with the overall pattern (Fig. 6) resembling the theoretical pattern of Fig. 5b. This suggests that the evolutionary differentiation between major taxa is constrained by a bound on mass specific metabolism, and that the body mass variation between the species of a given large bodied taxa is reecting mainly niche differentiation through the evolution of diverse resource handling. Note also that the transition from minimum sized organisms to larger organisms with 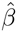 exponents around −1/2*d* fall, as predicted, between the largest unicells with asexual reproduction and the multicellular metazoan with sexual reproduction between a male and female individual.

**Figure 6:**
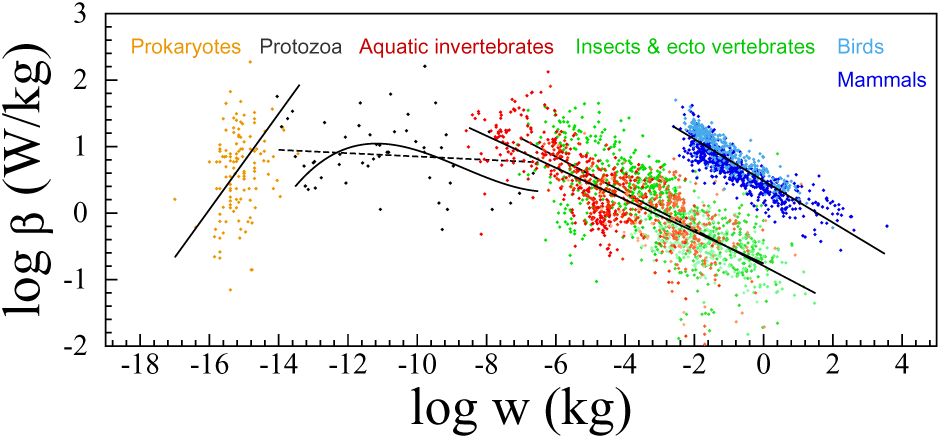
Heterotroph organisms. Macro evolutionary relationship between mass and minimum mass specific metabolism among heterotroph organisms. Data from Makarieva et al. (2008), with RMA lines from DeLong et al. (2010) for prokaryotes and protozoa, and least-squares lines for other taxa. Prokaryotes: 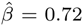, *n* = 123, for passive (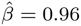 for active); Protozoa: 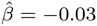, *n* = 52; Aquatic invertebrates: 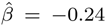, *n* = 808; Insects & ectotherm vertebrates: 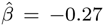, *n* = 982; Birds & mammals 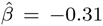, *n* = 948. A least-squares fitted third order polynomial is also shown for protozoa, excluding the left-top four species.

## 8 Discussion

By integrating primary selection on mass specific metabolism into the theory of Malthusian Relativity, I showed that the origin of replicating molecules can initialise a natural selection where the life history evolves as a balance between the energy that the average individual assimilates from the environment and the energy that it uses in metabolism and reproduction.

This balance is not just a physiological balance. The Darwinian paradigm implies evolution by differential self-replication. But it was found that the frequency independent selection of the physiology can select only for an initial transition from replicating molecules to self-replicating cells with a minimum mass. A balance with individuals that are larger than prokaryotes, including larger unicells and multicellular animals with sexual reproduction, larger unicells and multicellular animals with sexual reproduction, of the density-frequency-dependent selection of the interactive competition that follows from the population growth of self-replication.

This evolutionary unfolding of population dynamic feed-back selection is a unidirectional development given suitable environmental conditions, and it follows from a sustained selection for an increase in the net energy that is available for self-replication. The evolution of the major lifeforms was therefore found to occur as a deterministic succession from replicating molecules, over prokaryote and protozoa like self-replicating cells, to the fully developed multicellular animal with sexual reproduction between females and males.

This evolution raises the question why a majority of the heterotroph organisms have remained at a unicellular level. The straightforward answer is that they have been unable, for some reason or the other, to be selected to an energetic level that can sustain the feed-back selection for the multicellular animal.

If all phylogenetic lineages of mobile organisms were free to evolve by unconstrained selection across ecological niches we would, at least in the long run, expect that they would evolve towards the multicellular animal with sexual reproduction. But the inter-specific interactions of the larger species can exclude the smaller species from essential resources. Owing to this competitive exclusion of species by inter-specific competition, we can expect a distribution of species with net energetic states that range from a possible minimum value to a maximum, where the maximum is increasing over evolutionary time. In other words, while the intraspeci
fic selection is unidirectional when other things are equal, we expect a full range of lifeforms to be evolutionarily maintained by the competitive exclusion of inter-specific competition.

### 8.1 Theoretical background

During the development of the selection theory I integrated several concepts that were developed relatively independently of one another into the theory. Some of the essential components include *i*) metabolism as a proxy for the rate at which organisms assimilate, transform and expend energy (e.g. Calder 1984; Brown et al. 2004; Humphries and McCann 2014); *ii*) a biological time-scale as the inverse of mass specific metabolism (e.g. Pearl 1928; Brody 1945); *iii*) that advanced metabolism is dependent upon a cell where the molecules of the metabolic pathways can concentrate (e.g. Oparin 1957; Miller and Orgel 1974; Maynard Smith and Szathmáary 1995); *iv*) that natural selection is driven by the biochemical energetics of self-replication (e.g. Lotka 1922; Odum and Pinkerton 1955; Van Valen 1976; Brown et al. 1993); *v*) that it is constrained by physiological trade-offs and constraints (e.g. Charlesworth 1980; Roff 1992; Stearns 1992), including a metabolism that depends on mass in self-replicators with almost no mass (DeLong et al. 2010); *vi*) that it proceeds towards attractors like Continuously Stable Strategies (e.g. Maynard Smith and Price 1973; Eshel and Motro 1981; Taylor 1989); *vii*) that it is dependent upon the feed-back ecology of density dependence (e.g. Anderson 1971; Heino et al. 1998; Rankin 2007), including the density dependence of interactive competition (e.g. Abrams and Matsuda 1994; Witting 1997) that makes arms race models (e.g. Dawkins and Krebs 1979; Parker 1979; Maynard Smith and Brown 1986) realistic; *viii*) that the resulting short-term evolution is contingent upon the current state of biology and the available mutations; and *ix*) that long-term evolution is more like a deterministic path (Witting 1997, 2008) that is laid down by the selection attractors that unfold from the origin of replicating molecules, including allometric exponents that evolve by the ecological geometry of optimal density regulation (Witting 1995).

That an essential part of the long-term evolution of life histories is deterministic from the origin of replicating molecules is in contrasts to the 20th Century paradigm, where evolution by natural selection was seen as inherently contingent (e.g., Mayr 1988; Salthe 1989; Maynard Smith and Szathmáary 1995; Gould 2002; criticized by Kauffman 1993, 1995; Witting 1997, 2008; Conway-Morris 2003). In the strong interpretation, contingency implies an evolution that cannot be predicted *a priori* by natural selection, but can only be understood *a posteriori* from its historical development once it has actually occurred (Gould 2002). This view underlies classical life history theory that developed contingent *a posteriori* selection models (reviewed by Charlesworth 1980; Roff 1992; Stearns 1992) that explain traits from evolutionary assumptions on other traits in existing lifeforms.

The contingent view is typically seeing each population as evolving towards an optimum for the niche that it is occupying, with no inherent direction to natural selection *per se* (e.g., Stanley 1973; Gould 1988; Bonner 2006; Brown and Sibly 2006). An evolutionary size increase was seen as an adaptive advantage that may include an improved ability to capture prey, avoid predators, have greater reproductive success, decreased annual mortality, increased longevity, increased intelligence from increased brain size, expanded niche width, increased heat retention, etc. While these and many other factors may inuence the evolution of mass, the view of an adapting mass is insufficient in the sense that it does not develop explicit models on natural selection of mass.

Explicit models of contingency explained large body masses (McLaren 1966; Roff 1981; Stearns and Koella 1986), and allometric correlations (Kozłowski and Weiner 1997; Kozlowski et al. 2003b), from a reproductive rate that is positively correlated with mass across the body mass variation in a species. As reproduction is proportionally related to mass in many species (Peters 1983; Reiss 1989) these models mimic natural selection on mass in those species. But the view is insufficient in the sense that it does not explain the origin of the natural selection pressure on mass. The proportional dependence of reproduction on mass is instead explained by the deterministic model in this paper, where it evolves as a consequence of a resource bias of interactive competition in populations with body masses that are selected to increase exponentially (Witting 1997, 2003).

With Witting (2016a) and this paper I aimed for a natural selection that is sufficient to explain the joint evolution of the metabolism, mass, allometries, and major transitions that define lifeforms from viruses over prokaryotes and larger unicells to multicellular animals.

The proposed model is a sufficient Darwinian hypothesis that aims to explain the complete life history of the model organism from the origin of replicating molecules (Witting 1997, 2008). It is not a model that attempts to explain the evolution of everything, but merely a model that is internally self-consistent with the hypothesis that advanced lifeforms evolve as a deterministic consequence of the natural selection that unfolds from the origin of replicating molecules. This implies predictions from first principles of replication, without the estimation of parameters from data on evolved lifeforms.

My earlier work on sufficient population dynamic feed-back selection predicted an exponential increase in net energy and mass (Witting 1997, 2003), with a major transition between negligible-sized low-energy self-replicators with asexual reproduction, and high-energy organisms with large body masses, senescence, and sexual reproduction between a female and male individual; including additional transitions to cooperative breeding and eusocial colonies (Witting 1997, 2002a, 2007, 2008). The life histories of the larger species were predicted to include Kleiber scaling, with 1/4 exponents evolving from 2D interactions and 1/6 exponents from 3D (Witting 1995). This work followed a widespread tradition where the inter-specific variation in mass specific metabolism was seen as a consequence of the negative scaling with mass.

### 8.2 Pre-mass selection on metabolism

In Witting (2016a) and this paper I adjusted my first view on the natural selection of metabolism (Witting 2003), and separated the resource assimilation parameter of my original model into resource handling and the pace of handling. This pace generates gross energy, and by defining net energy as the difference between gross energy and the total metabolism of the organism, Witting (2016a) found the mass specific work of handling to be selected as mass specific metabolism. This implies primary selection for an increase in mass specific metabolism, an increase that generates the net energy that is a pre-condition for the selection of mass and the associated secondary rescaling of mass specific metabolism with the evolutionary changes in mass.

In the earlier version of my theory where metabolism evolved only by mass-rescaling, small bodied asexual self-replicators were predicted to be a single group with no explicit allometries. But by integrating primary selection on metabolism into the theory, the predicted asexual group unfolded into three sub-groups with life histories and allometries that resemble those of respectively viruses, prokaryotes and larger unicells. For these it is the dependence of mass specific metabolism on the minimum mass that is required to sustain metabolism that is the primary driver of selection; with transitions between the three lifeforms being selected from transitions in this dependence.

Replicating molecules with no intrinsic metabolism Replicating molecules with no intrinsic metabolism evolve when the maximum dependence of mass specific metabolism on minimum mass is so low that the increase in net energy from an initial increase in mass is too small to outbalance the energy that is required by the quality-quantity trade-off.

When the pre-mass exponent of this maximum dependence is larger than one, we predict prokaryote-like self-replicating cells with somewhat underdeveloped metabolic pathways. They evolve by a mass specific metabolism and mass that are selected as a balance between the quality-quantity trade-off and the dependence of mass specific metabolism on minimum mass. Unicells, that are larger than prokaryotes and with more developed metabolic pathways, are predicted to evolve when this selection balance is assisted by a population dynamic feed-back selection that has unfolded to some degree. This unfolding generates an increase in the interactive selection for mass, with the dependence of mass specific metabolism on minimum mass vanishing with an evolutionary increase in mass.

The point of evolution where the metabolic pathways are fully developed, and there no longer is a dependence of mass specific metabolism on minimum mass, was shown to coincide with the emergence of the fully developed population dynamic feed-back. The interactive competition is then selecting for masses are that larger than the mass that is required to sustain the rate of mass specific metabolism, and for a transition to the multicellular animal with senescence and sexual reproduction. Primary selection on metabolism will generate net energy for the selection of mass also in multicellular animals; yet the widespread Kleiber scaling in this group is predicted to reect variation in the handling and densities of the underlying resources.

### 8.3 The metabolic cell

The dependence of mass specific metabolism on mass in small replicators should be seen in relation to the evolution of the cell. Extremely low levels of metabolism are not in principle dependent upon the development of a compartment like a cell. But it is difficult to imagine the evolution of an advanced intrinsic metabolism unless it is connected with the formation of a cell, where the different molecules of the metabolic pathways can concentrate (e.g., Oparin 1957; Miller and Orgel 1974; Maynard Smith and Szathmáary 1995; Michod 1999; Wächtershäuser 2000; Koch and Silver 2005). Taking this point of view, I found that the increase in net energy for self-replication with increased metabolism is the primary selection that drives the evolution of the cell and all its associated mass. This implies that a self-replicating cell with heredity and an internal metabolism can be selected only from the subsets of the potential biochemistry that have a maximum pre-mass exponent for the dependence of mass specific metabolism on mass that is larger than unity (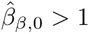).

This result is in agreement with the evolutionary emergence of the cell, as well as the evolutionary transition between unicellular self-replicators and multicellular organisms. With the cell being the compartment that organises metabolic pathways, it follows that the quality-quantity trade-off is selecting for virus-like replicators with no metabolism and the absence of a cell when 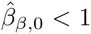. Unicellular self-replicators may instead evolve when 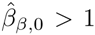, and as these are selected to have the minimum possible mass that is required for their metabolism, they are, as a consequence of this, selected to have no more than a single cell only. High-energy organisms with a sexual reproduction that is selected by interactive competition are instead predicted to have a mass that is larger than the required minimum for metabolism. Hence, they are free to evolve multicellularity with specialised and co-operating cells that enhance the behavioural performance of the individual and allow for the evolution of more efficient transportation networks within the organism.

## 9 Conclusion

The proposed natural selection is so far unique in the sense that it is the only one that is sufficient to explain the joint evolution of metabolism, mass, major life history transitions, and allometric exponents from viruses to multicellular animals. These transitions and exponents are explained almost as parsimoniously as possible as they evolve, not from separate processes, but from the primary selection on metabolism and mass itself; the selection that is necessary for the very existence of the organisms that are a pre-condition for the existence of life history transitions and allometries.

These organisms have to balance their energy between metabolism, mass and self-replication. The observed balance is only one of infinitely many potential possibilities, and the selection that unfolds from replicating molecules was found in this paper to explain the overall balance from viruses over prokaryotes and larger unicells to multicellular animals.

### 9.1 Eleven conclusions

1. Mass specific metabolism is predicted to be selected as the pace of the resource handling that generates net energy for self-replication, with persistent selection for a continued increase in net energy and mass specific metabolism.
2. The quality-quantity trade-off, where parents can produce many small or a few large offspring, is constantly selecting for the near absence of mass. This implies that the intra-specific dependence of net energy on mass needs to increase at least linearly with mass in order to outbalance the quality-quantity trade-off and select for larger body masses.
3. A selection increase in mass generates a mass-rescaling, where life history and ecological traits evolve in response to the evolutionary changes in mass. This mass-rescaling selection is initiated by a metabolic trade-off that selects for a dilation of the reproductive period with mass, in order to prevent a decline in net energy and fitness on the per generation time-scale of natural selection.
4. The physiological mass-rescaling of the life history imposes a mass-rescaling of the interactive foraging behaviour that determines the selection optimum of density regulation. And this ecological optimum explains the 1/4 exponents of Kleiber scaling as the two dimensional case of the more general 1/2*d*, where *d* is the spatial dimensionality of the intraspecific interactive behaviour.
5. Body mass allometries are predicted to depend also on a metabolic-rescaling, where the life history is rescaled by the relative importance of mass specific metabolism for the evolution of mass. 1/2*d* exponents evolve when the inter-specific body mass variation is invariant with respect to pre-mass variation in mass specific metabolism, and the corresponding exponents are (2*d* − 1)/2*d* when all of the variation in mass is evolving from pre-mass variation in mass specific metabolism.
6. Small self-replicators have a mass specific metabolism that is dependent also on the mass of the molecules in metabolic pathways, and on the mass of the cell where the metabolic molecules concentrate; including the mass of the heritable code for the cell and the metabolic pathways.
7. Replicating molecules with no metabolism and practically no mass - like viruses - are predicted to be selected from lineages where an initial mass specific metabolism is increasing sub-linearly with mass.
8. Small single-celled self-replicators with an internal metabolism are selected from lineages where the initial mass specific metabolism is increasing stronger than linearly with mass. Given the absence of an intra-specific resource bias from interactive competition, these self-replicating cells are predicted to have asexual reproduction and a mass specific metabolism that increases to the 5/6 power of mass (3D ecology), as empirically estimated in prokaryotes.
9. Larger single-celled self-replicators with a more developed metabolism and asexual reproduction are selected from a population dynamic feed-back of interactive competition that is unfolding gradually from an expected evolutionary increase in net energy. These interacting self-replicators have an allometric exponent for mass specific metabolism that declines from 5/6 over zero to −1/6 with an increasing mass (3D ecology), as empirically indicated for protozoa.
10. Multicellular animals with large body masses and sexual reproduction are predicted to be selected from the intra-specific interactive competition in high-energy lineages with a fully developed population dynamic feed-back selection. Given a species distribution that diversity across ecological niches, the predicted inter-specific exponent for mass specific metabolism is −1/2*d*, with the −1/4 ↔ −1/6 transition being observed quite commonly between terrestrial and pelagic taxa of multicellular animals.
11. Given a mass specific metabolism that evolves around an upper bound, mass specific metabolism is predicted to be body mass invariant on the macro evolutionary scale from prokaryotes to mammals, as empirically estimated by the absence of an allometric correlation.

## Appendix

## A Physiological selection on mass

This appendix describes some general considerations and definitions in relation to the physiological selection that is acting constantly on mass. Apart from a dependence of mass specific metabolism on mass in self-replicators with minimum body masses (see Section 3), I will treat the life history as being functionally independent of mass.

While this assumption will work in relation to the long-term evolution of mass, there can be special cases where life history components are functionally dependent on mass. Larger individuals may, e.g., be less likely to be taken by predators. This may inuence the distribution of evolved body masses, but it will generally not explain the evolution of large masses. Not only does it not apply to top-predators, but to explain the evolution of mass it requires that the survival parameter *p* increases at least proportional with mass, which is implausible, at least in general.

Another apparent link is net energy (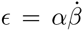) that nearly always is strongly positively correlated with mass. However, there seems to be no general mechanism that will explain the evolution of large body masses from a resource assimilation that is larger in larger organisms because resource assimilation necessarily is functionally dependent on mass. So I treat resource assimilation as functionally independent of mass, aiming to show that large masses evolve by natural selection in high energy species (high energy organisms could instead have evolved small masses and very high replication rates).

For this I assume that it is generally not a functional dependence of *α* on *ω* that is the cause for the evolution of large body masses. Nevertheless, the handling of resources may indeed increase mechanistically with mass. But this dependency is assumed to be so small that it will generally not affect our evolutionary predictions if it is ignored. To show that large species can evolve by natural selection we need to predict that they evolve a strong positive dependence of *ω* on *α* (i.e., if *β*_*β*_ is constant); and with *α* being functionally independent of *ω* we predict the evolution of a mass that often increases proportional with *α* (Witting 2016a). This is a stronger link than we would expect for almost any functional dependence of *α* on *ω*.

Maybe the strongest functional dependence of *α* on *ω* is in organisms that obtain resources through their complete surface. In this case we may expect *α* ∝ *ω*^2/3^ from the relationship between the surface area and the volume (mass) of an individual. Yet, from eqns 1 and 2 we find that *α* must increase at least proportional with mass for a large body mass to evolve as a response to a positive dependence of *α* on *ω*. Hence, even in this special case with a very strong dependence, we cannot straightforwardly explain large masses as a selection consequence of a positive linking of *α* to *ω*.

The availability of resources *ρ*, which is included in the *α* parameter (as 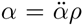, where 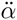 is intrinsic handling), might also increase with mass if an increased mass allows the organisms to exploit a larger spectrum of resources. While such a positive relationship may exists in some cases, and while this may inuence the distribution of evolved body masses, it is unlikely a general principle that will explain the evolution of large masses. Quite generally, I treat resource availability as a parameter that is not directly part of the phenotype and thus not affected directly by the natural selection of my model (but it may be affected indirectly by the evolution of the phenotype). Nevertheless, variation in *ρ*^**^ is affecting the realised net energy, and this variation is selected into variation in mass; as part of the variation in *α* that is selected into variation in mass.

As none of these potential links of the life history to mass will explain a general selection for large body masses, I will usually treat the life history as being independent of mass except for the dependence that evolves from the natural selection of my equations. The selection gradient on log body mass that is imposed by the physiology is thus minus one

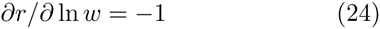

from eqn 1, and this reects a replication cost of mass that is selecting for a minimum mass

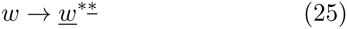

I treat this downward pull as a fundamental background that any natural selection for mass needs to outbalance in order to push evolution beyond the replicating molecules at the origin of life.

### A.1 Classical selection theory

The downward pull of eqn 24 was neglected in the contingent selection of classical life history theory (McLaren 1966; Roff 1981, 1986; Stearns and Crandall 1981; Stearns and Koella 1986). The contingency made it possible to assume an intra-specific relation between mass and replication that was based on observations instead of energetic trade-offs. The reproductive rate was thus assumed to be about proportional to mass, as observed in many populations (Peters 1983; Reiss 1989) and supported by widespread natural selection for an increased mass in natural species (Kingsolver and Pfennig 2004).

To describe this by a simple contingent model, we note from eqn 1 that intra-specific proportionality between reproduction and mass requires net energy to increase to the second power of mass (ε ∝ *ω*^2^). Then, if survival is negatively related to mass (e.g., as *ρ* ∝ *e*^−*cw*^ we may obtain a fitness profile (landscape) *r*_*i*_ = ln *ω*_*i*_ − *cw*_*i*_ that describes fitness as a function of the body mass variation (*ω*_*i*_) in the population (Fig. 1a). The differential of the profile 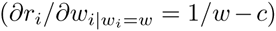 at the limit of the average variant (*ω*) is the selection gradient (Fig. 1b), that describes selection as a function of the average body mass. And the evolutionary equilibrium (*ω*^**^ = 1/*c*) is the optimum of the selection integral [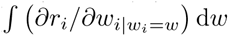; Fig. 1c], that integrates the selection gradient across the potential evolution in the average mass of the population.

The more recent approach of metabolic ecology (Brown and Sibly 2006) assumes, somewhat contrary to evidence, that the variation in the intra-specific birth rate follows the inter-specific allometry for biotic periods (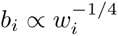), with an intra-specific birth rate that declines with mass instead of increasing as often observed. Large species may then evolve by a survival rate that increases with mass. While metabolic ecology have changed the selection roles of reproduction and survival relative to traditional life history theory, the two approaches are structurally identical as they are based on constant relative fitnesses with frequency independent selection. The outline in Figs. 1a, b and c is therefore applicable to both, as well as to other arguments and models with frequency independent fitness on mass (e.g., Schoener 1969; Stanley 1973; Charlesworth 1994; Charnov 2011; DeLong 2012).

The plots in Figs. 1a, b and c show a direct relationship between the fitness landscape (profile) and the potential evolution along the selection integral, with the most fit variant in any population being the average variant at the evolutionary equilibrium. This relation reects evolution by Fisher’s (1930) fundamental theorem that is selecting for an increase in what is known as the average fitness of the population; measured either by the intrinsic growth rate (*r*) and/or by the carrying capacity (*n*^*^) of the population (Roughgarden 1971; Charlesworth 1971; Witting 2000a).

One inherent problem with these approaches are the constant relative fitnesses that make the intrinsic components of fitness, and of fitness related traits like reproduction and survival, scale identically within and across species (Witting 2000a, 2002b); as illustrated by the similar curves in Figs. 1a and c. This restriction is quite generally contradicted by data in at least birds and mammals, where lifetime reproduction and survival tend to be independent of mass across species (Witting 1995).

Also, for inter-specific allometries in multicellular species, it is well-known that *r* tends to scale to the negative 1/4 power of mass (Fenchel 1974) and that *n*^*^ tends to scale the negative 3/4 power (Damuth 1981). In combination with a life history theory that is based on constant relative fitnesses, these data imply a decline in fitness (*r* or *n*^*^) with mass, and this generates the paradox of constant selection for the near absence of mass in basically all natural species. Realized reproduction and survival could, of course, scale differently at the two scales if resource exploitation was a function of body mass across species. But the resource consumptions of populations are often independent of mass across natural species (Damuth 1981; Peters 1983; Calder 1984).

We may thus conclude that the combination of contingency and frequency independent selection is insuff-cient to explain the joint evolution of mass and allometries.

